# Genome-wide mapping of DCP2-dependent 5′ cap footprints in *Arabidopsis thaliana*

**DOI:** 10.64898/2026.03.04.709597

**Authors:** Neha Shukla, Michael A. Schon, Vivek K. Raxwal, Michael D. Nodine, Karel Riha

## Abstract

mRNA decapping mediated by DCP2 is a key mechanism controlling RNA stability and gene expression in eukaryotes, including plants. Despite its central role in regulating plant development and stress responses, the repertoire of mRNA 5’ caps targeted by DCP2 remains undefined. Here, we combined *in vitro* decapping treatment with 5’-end enriched and full-length transcriptome sequencing of DCP2-deficient mutants to comprehensively characterize the mRNA capping landscape in *Arabidopsis thaliana*. We mapped over 13,000 high-confidence capped transcripts at nucleotide resolution, revealing distinct 5’ cap signatures in both wild type and *dcp2* seedlings. Most caps were localized near annotated transcription start sites, validating the accuracy of our approach. Loss of DCP2 led to a substantial accumulation of capped mRNAs, including 275 capped transcripts originating from previously unannotated loci. It also increased prevalence of multi-capped genes highlighting the role of DCP2-mediated decapping in removing unwanted transcripts. Integration of these data with degradome resources revealed that targets of co-translational and cytosolic XRN4-dependent decay, as well as of nonsense-mediated decay, were enriched among capped mRNAs specifically accumulated in *dcp2*. These findings suggest that mRNA degradation in these decay pathways is mediated through decapping. In addition, this study provides a valuable resource for transcript annotation and isoform-aware analysis of RNA turnover.

**GRAPHICAL ABSTRACT:** 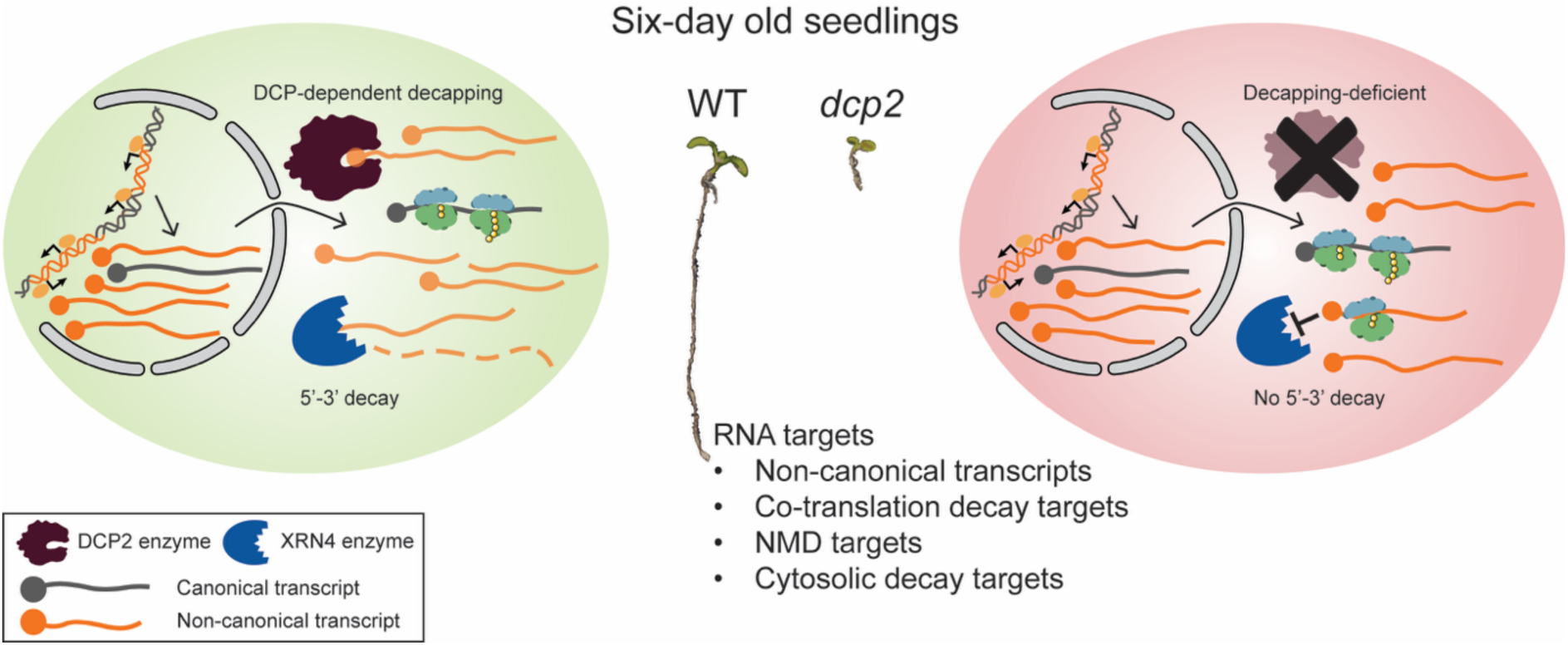

## INTRODUCTION

Gene expression is largely determined by the steady-state levels of mature mRNA, which arise from balance between RNA synthesis and decay (1, 2). mRNA decay is not a random process but proceeds through specific, well-defined pathways (3–5). In the predominant 5’-3’ decay, deadenylation is followed by removal of the protective 5’ cap (m^7^GpppN) through decapping and subsequent exonucleolytic degradation by ribonuclease XRN1 or its plant homologue XRN4 (4, 6–8). Decapping is responsible for the turnover of bulk cytoplasmic mRNA (9–11), as well as transcripts targeted by specialized surveillance mechanisms, including nonsense-mediated decay (NMD) (12–14), microRNA-mediated decay (15, 16), degradation of AU-rich elements (ARE) -containing mRNAs (17), and transcripts with non-optimal codons (3, 18). Thus, decapping represents a key step in mRNA turnover, making it central to the post-transcriptional regulation of gene expression.

The cleavage of 5’ caps is mediated by the DCP1/2 decapping complex (19). The primary catalytic subunit of this complex is DCP2, whose activity is enhanced by DCP1 (20–23). DCP2 is a member of the Nudix (Nucleoside Diphosphates linked to moiety X) hydrolase family (24–26). Among numerous nudix hydrolases, DCP2 exhibits selective activity toward m^7^G-capped mRNAs by forming a cytoplasmic multi-protein complex with other decay factors (27–31).

Decapping has been very well defined across all eukaryotic organisms. Decapping-deficient mutants display pleiotropic developmental and stress-responsive phenotypes, underscoring the essential role of decapping in normal physiological processes. The compromised decapping in *Caenorhabditis elegans* and *Drosophila melanogaster* results in impaired growth and development, shortened lifespan, and increased stress sensitivity (32–34). In mice, defective decapping results in slow growth phenotype and enhanced innate immune responses, while the postnatal depletion of DCP2 leads to male infertility (17, 35). In plants, the core decapping complex consists of DCP1, DCP2 and scaffold Varicose (VCS), and Arabidopsis mutants lacking any of these genes exhibit seedling lethality (36, 37). In addition to its role in development, decapping is important for the precise degradation of specific transcripts to ensure proper regulation of various cellular processes, especially during stress responses (6, 17, 38–46).

Analysis of RNA decay rates in Arabidopsis *vcs-7* null mutants suggests that decapping contributes to the decay of up to 68% of the transcriptome (10). However, *in vivo* substrates of DCP2 and the precise positions of canonical 5’ caps, have not been comprehensively mapped. Here, we use low input mRNA 5’-end enriched sequencing (47), in combination with *in vitro* decapping, to accurately delineate 5’ cap footprints of mRNAs in Arabidopsis seedlings. Comparison of wild type and *dcp2* mutants revealed transcripts subjected to decapping-mediated turnover, thus providing an important resource for understanding the role of mRNA decay in transcriptome homeostasis.

## MATERIAL AND METHODS

### Plant materials and growth conditions

*Arabidopsis thaliana* Columbia-0 (Col-0) ecotype was used as wild type. The T-DNA insertion mutants, *dcp2-1* (SALK000519), *dcp1-1* (GK-844B03.02), *vcs-6* (SAIL_831_D08), *vcs-7* (SALK032031) and *eds1-2* (48) that were used in this study were obtained from NASC. Seeds were sterilized and plated on agar plates containing half-strength Murashige and Skoog (MS) medium, adjusted to pH 5.7, supplemented with 1% (w/v) sucrose. Plates were cultivated in growth chambers under 16/8-hour light/dark cycle at 22°C. All the decapping-deficient single or double mutants used in the study were PCR genotyped to confirm the presence of T-DNA insertions (Supplementary Table S1).

### *In vitro* decapping assay, library construction and sequencing

For RNA isolation, six-day old seedlings of wild type and *dcp2-1* homozygous mutants were harvested. Seedlings from each plate were pooled and considered one biological replicate. For each genotype, total RNA was isolated from two independent biological replicates using RNA Blue (TOP-Bio, Catalog No. R013). Following RNA extraction, samples were treated with TURBO DNase (Thermo Fisher Scientific, Catalog No. AM2238) to remove genomic DNA contamination. The RNA was then purified using magnetic beads (Beckman Coulter, Catalog No. A63987), following the manufacturer’s instructions. Total RNA from each sample was split into two parts, one treated with DCP2 enzyme and the other with no treatment.

For *in vitro* decapping assay, 5 μg total RNA samples were treated with the mRNA Decapping Enzyme (NEB, Catalog No. M0608S) at 37 degrees Celsius for 30 minutes. RNA was purified post-treatment using magnetic beads (Beckman Coulter, Catalog No. A63987) to ensure clean separation from the enzyme and other reaction components. Then, we used a modified low-input Parallel Analysis of RNA Ends (nanoPARE) protocol as described by Schon et al., 2018 (47) for library preparation. For cDNA synthesis, 1 μg treated RNA or 1 μg non-treated RNA was reverse-transcribed with anchored oligo-dT primer and TSO oligo using SuperScript IV reverse transcriptase (Thermo Fisher Scientific, Catalog No. 18090050) using recommended cycle conditions. cDNA synthesis was followed by pre-amplification and was purified using SPRISelect magnetic beads (Beckman Coulter, Catalog No. B23318). The integrity and quality of pre-amplified cDNA was assessed by a fragment analyzer (CF Genomics, CEITEC). Following pre-amplification, 1 ng pre-amplified cDNA was used to prepare both 5’-end enriched libraries and full-length whole transcriptome libraries. Briefly, cDNA was tagmented and library amplification was performed using NextraXT kit (Illumina, Catalog No. FC-131-1024) following manufacturer’s protocol.

The final libraries were checked on a fragment analyzer and subsequently sequenced. Transcriptome libraries were sequenced 75 bp single-end using standard Illumina sequencing primers and 5’-end enriched libraries were sequenced 150-bp single-end using a mix of custom sequencing primers on Illumina NextSeq platform at CEITEC genomics core facility.

### Data processing and analysis of transcriptome reads

The low-quality reads and adapters were first trimmed from the total reads using cutadapt v4.1 (49). Then, the RNA-Seq data was aligned to the reference AtRTD3 transcript dataset (50) and simultaneously quantified using Kallisto software v0.50.1 (51). The downstream data normalization and differential expression analysis was done using the 3D RNA-Seq tool (52, 53).

In brief, the read counts and transcripts per million reads (TPM) were generated and normalized to transcript length over the samples and the library size using tximport R package (version 1.10.0) and lengthScaledTPM method. The low expressed genes were filtered based on data mean-variance trend. A gene was expressed if it has at least one expressed transcript with 1 count per million mapped reads (CPM) in two or more samples. The trimmed mean of M-values (TMM) method was used to normalize the gene read counts to CPM. The voom-pipeline of limma R package was used for pair-wise comparisons of gene expression between the samples. Statistical analyses were performed using Student’s t-test or the Wilcoxon rank-sum test, as indicated in the figure panel, to evaluate differences in expression between the samples. Corresponding p-values are reported in the figures. A gene was significantly differentially expressed (DE) in a sample if it had adjusted p-value < 0.05 (Benjamini-Hochberg (BH)-corrected) and absolute log_2_FC ≥ 1. The previously published RNAseq dataset for wild type and *vcs-7* (GSE86359)(10) was reanalyzed in a similar way for comparisons with present study.

### Data processing and analysis of nanoPARE reads

Sequencing data from six-day-old seedlings generated in this study have been deposited in the Gene Expression Omnibus (GEO) under accession number GSE315704. For each sample, 5’ caps were identified using the nanoPARE analysis pipeline (47). Briefly, EndMap module removes the adapter sequences and filters low complexity reads using cutadapt (v4.1). Then it aligns the reads to TAIR10 reference using STAR aligner (v2.7.1) with default parameters for each sample. After alignment, it applies sequence-bias correction (54) (adapted for RNA), computes per-base coverage of uniquely mapped reads and assigns multi-mapping reads with a “rich-get-richer” scheme. The module outputs strand-specific coverage of 5’ end counts across the genome and for 5’-end enriched libraries, it records 5’ soft-clipped bases as upstream untemplated nucleotides (uuNs). Also, the known artifacts (TSO mispriming and internal oligo-dT) were masked. Further, EndGraph module evaluates each sample pair (5’-end enriched with corresponding transcriptome) to predict 5’ features per replicate with a peak at the local maxima via subtractive kernel density estimation. Briefly, to reduce expression-dependent background, the 5’ ends were normalized with corresponding transcriptome (single scale per sample). This signal was smoothed by kernel density estimation and genome-wide discrete 5’ features (cut-off = RPM ≥ 0.5) were identified. Next, EndClass module was employed where replicates of the same sample type were merged to re-identify the peaks and only reproducible features were retained. Features were assigned to nearby transcripts (including ≤500 nt upstream of 5’-terminal exons) and labeled by exon context. Later, they were classified as capped or non-capped based on the proportion of upstream untemplated guanosines (uuG) within each feature. In the present study, features with ≥5% uuG are considered capped.

### General data processing and analysis

Genomic overlaps with previously published data were performed using gene IDs, wherever available, to identify 5’ cap features of co-translational and cytosolic decay targets, nonsense-mediated decay (NMD) targets and immune receptor transcripts. In case of unannotated loci, overlaps was performed using genomic coordinates in R (v4.4.2) with bioconductor package Genomic Ranges (v1.58.0) (55). Shared 5’ features between samples were quantified using BED Tools intersect (v2.27.1) in a strand-specific manner (56, 57). All the graphs and plots were generated in R (v4.4.2) and the statistical significance of feature overlaps was assessed using Fisher’s exact test. The 5’ feature overlaps between wild type and *dcp2* samples were visualized using online interactive venn diagram tool (58). The 5’ cap enrichment index was calculated as a change in cap signal estimate. For each gene, we calculated the log_2_-fold change (*dcp2* vs wild type) independently using 5’-end RPM-based counts and full-length TPM-based expression. Then, the 5’ cap enrichment index was defined as the difference between these two values, indicating whether the 5’-end signal increases more than overall gene abundance.

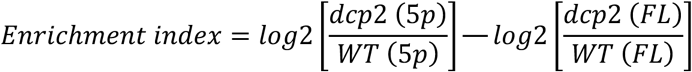

Where *5p* is the 5’-end expression, *FL* is the full-length transcript expression and *WT* is wild type sample.

The functional annotation of genes, their characteristics, and Gene Ontology (GO) enrichment analyses were performed online using the ShinyGO v0.81 tool (59). False discovery rate (FDR)-adjusted p-values were calculated based on the hypergeometric distribution, and GO terms were considered significant if the FDR was less than 0.05. For clarity and better visualization, the coverage plots were edited after export from Integrative Genomics Viewer (IGV) browser (60).

## RESULTS

### Transcriptome analysis in DCP2-deficient plants

To assess the impact of decapping on the Arabidopsis transcriptome, we analyzed *dcp2* null mutants. Growth of *dcp2* mutants is arrested approximately six days post-germination at the cotyledon stage, leading to seedling death due to necrosis within 12-14 days post-germination (36). Seeds from plants heterozygous for *dcp2-1* germinated on agar plates produced a fraction of growth-arrested seedlings that were validated by PCR to be homozygous mutants. Given the lethal nature of *dcp2* mutants, we analyzed the transcriptome of six-day old seedlings, which despite growth irregularities, still exhibit green, pre-necrotic leaves (Fig. 1A). We used the SmartSeq2 method to construct the full-length poly(A) transcriptome libraries for wild type and *dcp2* mutants, each with two biological replicates.

**Figure 1:**
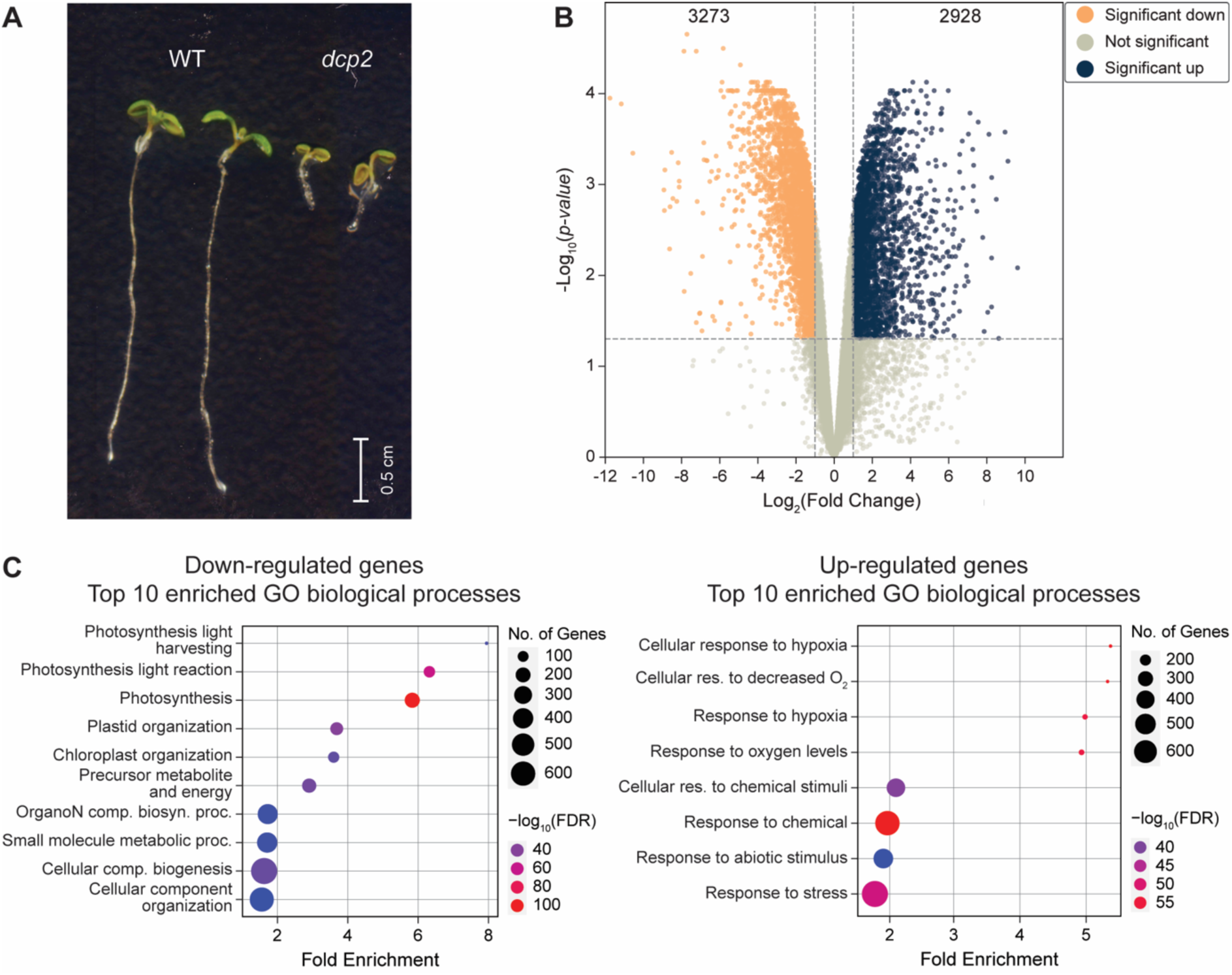
Loss of DCP2 remodels the seedling transcriptome. *(A)* Phenotypes of six-day-old seedlings of wild type and *dcp2* null mutant. *(B)* Volcano plot of differentially expressed genes (log_2_FC) identified in *dcp2* compared with wild type. The significantly altered genes are color coded; down-regulated genes are yellow and up-regulated genes are blue. *(C)* Gene Ontology enrichment analysis with dot plots showing top 10 enriched GO biological processes among up- and down-regulated genes in *dcp2* compared with wild type (adjusted p-value < 0.05). The most to least significantly enriched categories are colored as red to purple, and the size of circle correspond to the number of genes in the category. *FC*, fold change; *GO*, gene ontology; *WT*, wild type.

Transcriptome analysis revealed that compared with wild type, DCP2 deficiency significantly altered the expression of 6,201 genes (log_2_ fold change ≥ 1 or ≤ -1; adjusted p-value < 0.05), with 52.8% genes (3,273/6,201) showing decreased expression (Fig. 1B). The functional annotation using gene ontology (GO) enrichment analysis indicated that biological processes related to stress responses, including hypoxia and abiotic stimulus, were notably enriched among up-regulated genes (Fig. 1C). Conversely, processes such as carbon fixation, photosynthesis, primary amino acid and metabolite biosynthesis, organelle and cellular organization were significantly down-regulated in *dcp2* mutants (Fig. 1C). These signatures in gene expression are consistent with pre-senescence and growth arrest phenotype observed in the seedlings.

Similarly to *dcp2* plants, Arabidopsis null mutants in other components of the decapping complex also exhibit post-embryonic lethality (Supplementary Fig. S1A) (36). To assess whether deficiency in different decapping factors elicit similar response, we re-analyzed the previously published transcriptome of *vcs-7* (10) and compared it with that of *dcp2*. As expected, a moderate correlation (Pearson’s r = 0.62, n = 17,317) was observed between the *dcp2* and *vcs-7* transcriptomes (Supplementary Fig. S1B). The GO enrichment analysis confirmed common activation of stress responsive genes, including those involved in cellular response to hypoxia, and the suppression of genes required for generation of precursor metabolites and energy in both mutants (Supplementary Fig. S1C). These results suggest that as a fundamental process, decapping affects multiple cellular pathways.

### Mapping canonical 5’ mRNA caps

Next, we sought to complement the transcriptome analysis with a genome-wide mapping of the 5’ cap positions. To identify capped 5’ ends of mRNAs, we combined a low-input Parallel Analysis of RNA Ends (nanoPARE) sequencing (47) with *in vitro* decapping treatment using DCP2 enzyme for both wild type and *dcp2* mutant (Supplementary Fig. S2). Prior to cDNA library preparation, total RNA from each sample was split into two parts, one treated with DCP2 enzyme (T) and the other with no treatment (NT). Subsequently, two cDNA libraries were created from each DCP2 treated and untreated RNA, one for 5’-end enrichment (nanoPARE) and the other for full-length transcriptome (SmartSeq2). Two biological replicas of wild type and *dcp2* seedlings yielded a total 16 libraries, comprising approximately 400 million raw reads (Supplementary Table S2).

We employed the nanoPARE bioinformatics pipeline to identify 5’ ends bearing caps at single-nucleotide level resolution (47). The pipeline identifies 5’ caps by the presence of an untemplated upstream guanosine (uuG) at the junction between the genomic sequence and the template-switching oligonucleotide (TSO). Briefly, after adapter trimming and per-strand genome alignment using STAR (61), uniquely mapped reads were used to identify high-confidence 5’ ends as described in the Methods section. We defined 5′ features representing local clusters of 5’ ends and classified a feature as capped when reads containing uuG exceeded 5% of all 5′ reads within that feature. Features falling below this threshold were considered as non-capped. This approach enabled delineation of genomic 5’ cap sites while effectively managing artifacts and biases.

This analysis identified a total of 19,266 5’ features from the unified no treatment samples of wild type and *dcp2*. Among these, 15,797 (81.9%) were classified as capped and 3,469 (18.1%) were classified as non-capped features (Supplementary Table S3). In DCP2-treated samples, the proportion of capped 5’ features decreased from 82% to less than 5% of all 5’ features (Fig. 2A; Supplementary Tables S4<u>-S7</u>). We checked the effect of *in vitro* DCP2 treatment on overall transcript abundance and a significant correlation was observed in gene expression before and after treatment in both wild type (Pearson’s r = 0.97) and *dcp2* (Pearson’s r = 0.96) (Supplementary Fig. S3A). Also, no significantly differentially-expressed genes were identified among DCP2-treated vs no treatment samples in both wild type and *dcp2* (Supplementary Fig. S3B), indicating that overall transcript abundance was only slightly reduced. These results indicate that *in vitro* DCP2 treatment efficiently and non-selectively removes 5’ caps without substantially affecting total RNA abundance. Further examination of the distribution of 5’ features across different samples revealed 11,246 capped sites in wild type and 14,125 in *dcp2*, whereas DCP2 treatment reduced the number of cap sites to 580 and 237, respectively (Fig. 2B; Supplementary Tables S6<u>-S7</u>).

**Figure 2:**
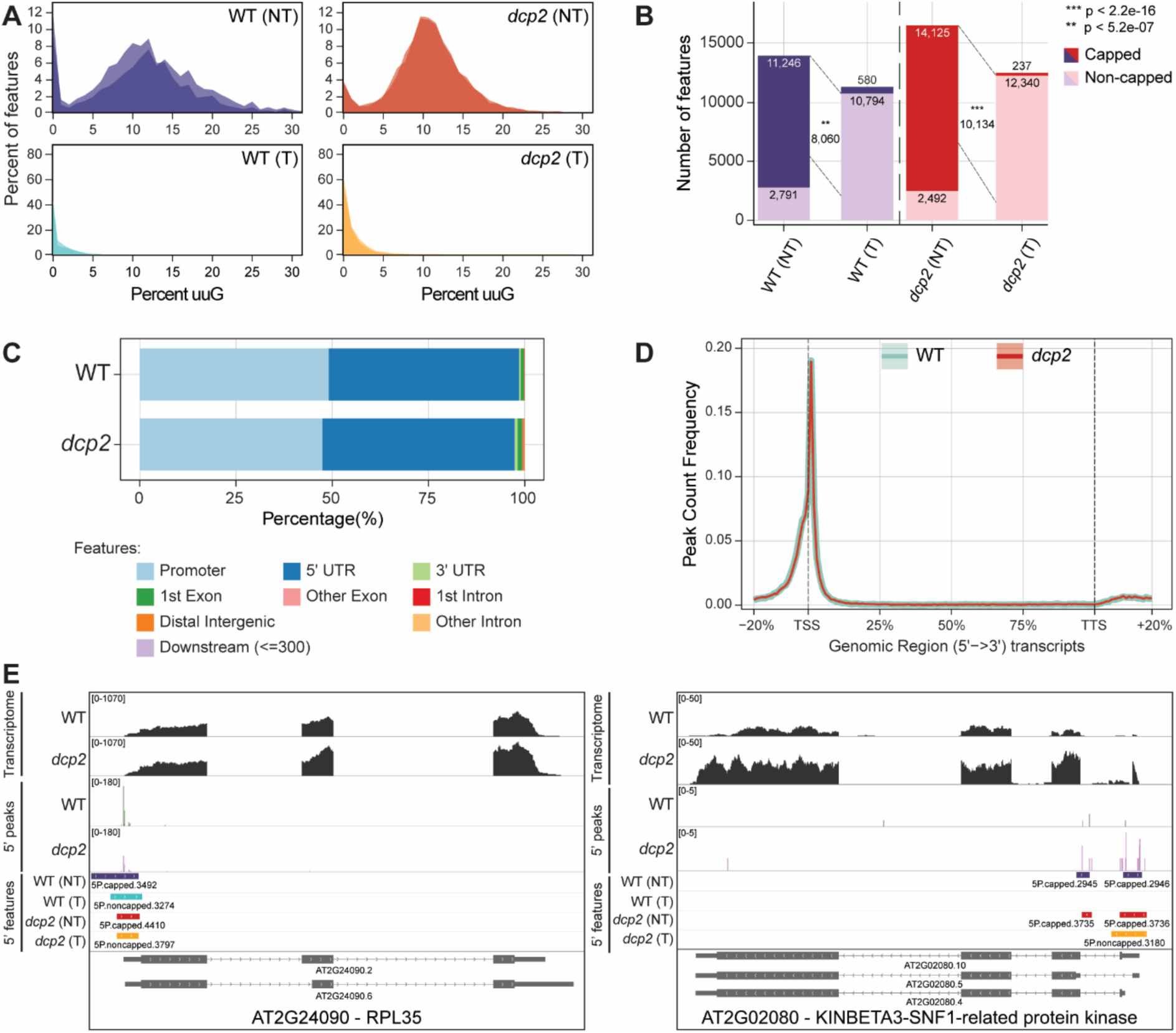
Identification of canonical 5’ features in Arabidopsis seedlings. *(A)* The distribution of 5’ ends represented as the proportion of 5’ features in nanoPARE reads (y-axis) with percentage of untemplated upstream guanosine (uuG; x-axis) from either no treatment or DCP2-treated fractions in both wild type and *dcp2*. *(B)* The number of 5’ features (y-axis) classified as either capped or non-capped, identified from wild type and *dcp2* (x-axis) from either no treatment or DCP2-treated fractions. The number of overlapping ends between the no treatment capped and treated non-capped in each group is indicated between the respective sample bars. The asterisk mark indicates the significance of overlap between the groups, determined by the Fisher’s exact test. *(C)* The distribution of identified 5’ caps over the genomic elements as annotated in AtRTD3 transcriptome. *(D)* The positional distribution of 5’ caps overlapping gene-body of protein-coding transcripts. *TSS*, Transcription start site, *TTS*, Transcription termination site. *(E)* IGV browser track showing the coverage (y-axis) of transcriptome and 5’ peaks with example of a ribosomal protein-encoding RPL35 gene (At2g24090) and SNF1-related protein kinase (At2g02080) identified in wild type and *dcp2*. The 5’ ends as identified in each sample as well as the gene structure of encoded transcripts [UTRs (narrow rectangle), exons (broad rectangle) and introns (lines connecting the exons)] from IGV are displayed. *IGV*, Integrated genome viewer; *NT*, No treatment; *T*, DCP2-treatment; *WT*, wild type.

In plants, mRNA turnover is primarily influenced by decapping followed by 5’-3’ exonucleolytic degradation mediated by the cytoplasmic ribonuclease XRN4 (62–66). To assess consistency with previous studies, we compare all 5’ features identified here with nanoPARE datasets generated from floral buds of wild type and *xrn4* mutants (47). This analysis revealed an 85.9% overlap (p < 2.2e-16 using fisher’s exact test) in detected 5’ features (Supplementary Fig. S3C), confirming reproducibility of this approach.

### Capped 5’ features are concentrated around transcription start sites

We next examined the genomic distribution of 5’ features. The majority of 5’ features were associated with protein-coding genes (10,456 (92.9%) in wild type; 13,082 (92.6%) in *dcp2*), aligning predominantly within annotated promoters or 5’ UTRs (Fig. 2C). The presence of a 5’ cap typically marks a transcription start site (TSS), which is generally located within a transcription start region encompassing several alternative TSSs. Alternative TSS usage contributes to transcript isoform diversity, representing an important mechanism influencing post-transcriptional regulation and protein function (67, 68). In this analysis, each 5’ feature can be considered equivalent to a transcription start region, while the 5’ cap position is the peak of uuG within a feature that correspond to a TSS. We assessed the positional distribution of 5’ caps in protein-coding genes by comparing their locations with transcript annotations from AtRTD3 (50). The majority of 5’ caps were located at or near annotated TSSs (Fig. 2D). Examination of individual genes confirmed the 5’ caps were typically positioned at the annotated start site (e.g. *RPL35*) or represented alternative capped positions within gene bodies (e.g. *SNF1*-related protein kinase) (Fig. 2E). Further comparison revealed that 5’ features were slightly broader in *dcp2* compared with the wild type, with a median width of 124 nt and 103 nt, respectively (Supplementary Fig. S3D). Broader 5’ features in *dcp2* may indicate a greater number of TSSs distributed across a wider span. This suggests that in wild type, decapping followed by rapid degradation may restrict the accumulation of a subset of transcripts originating from such alternative TSSs.

### Genome-wide landscape of 5’-end caps in *dcp2* mutants

Despite the efficient *in vitro* decapping reaction, a small number of residual caps were detected in DCP2-treated samples (Fig. 2B). These residual caps were excluded from further analyses, yielding a total of 10,846 high-confidence 5’ caps in wild type and 13,965 in *dcp2* (Supplementary Tables S8<u>-S9</u>). Of these, 9,488 (p < 2.2e-16 using fisher’s exact test) were shared between the two genotypes (Fig. 3A), indicating a stable core TSS landscape and strong assay concordance. The *dcp2* mutants exhibited a higher proportion of genotype-specific caps (4,488) compared with wild type (1,397) (Fig. 3A). These genotype-specific caps largely reflect a general increase in the expression of associated genes within the corresponding categories (Fig. 3B). In addition, we calculated 5’ cap enrichment indices to compare the abundance of 5’-end signal over full-length gene in *dcp2* relative to wild type. We used the 5’ cap enrichment index as a proxy to quantify changes in cap signal in absence of DCP2 as described in Methods section. The strongest 5’ cap enrichment index was observed for *dcp2*-specific capped genes (n = 4,153; Fig.S3E), indicating that these genes show significant increase in capped 5’-end abundance in *dcp2*. Shared capped genes (n = 9,263) also displayed a positive enrichment index (Fig. S3E), suggesting that many capped RNAs detected in both genotypes exhibit a stronger cap signal in *dcp2* compared with wild type. In contrast, WT-specific caps (n = 1,369) showed the weakest 5’ cap enrichment, implying no or small increases in cap accumulation in *dcp2* (Supplementary Fig. S3E).

**Figure 3:**
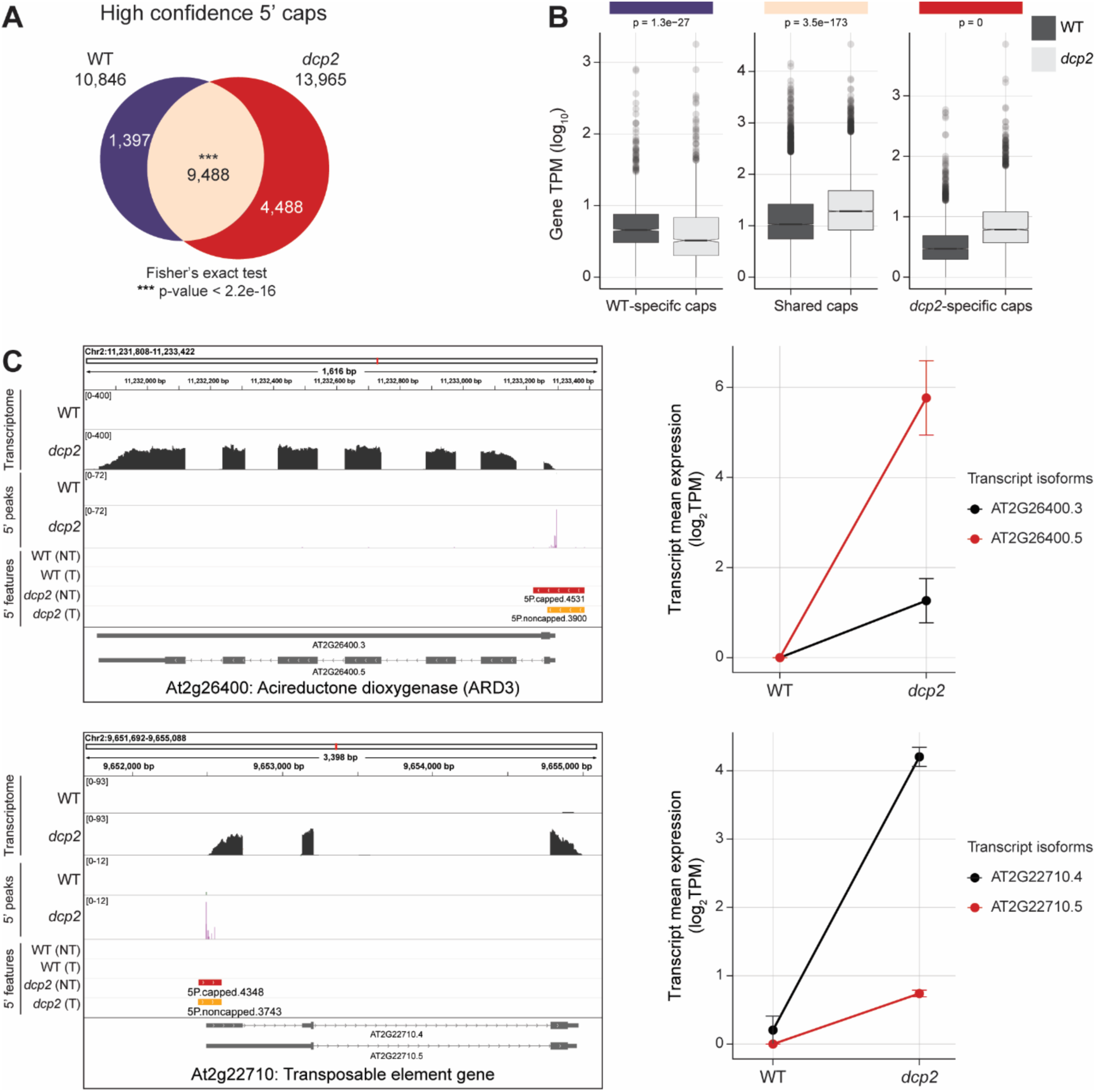
Genome-wide landscape of 5’ end caps in *dcp2* mutants. *(A)* Venn diagram showing the comparison between high-confidence 5’ caps identified in wild type and *dcp2*. The overlapping region (beige) are the caps shared between the samples while the non-overlapping regions depict 5’ caps unique to either wild type (purple) or *dcp2* (red). *(B)* Boxplot showing the expression (log_10_TPM) of genes that are wild type-specific capped genes (1,397; in purple), shared capped genes (9,488; in beige) or *dcp2*-specific capped genes (4,488; in red). *(C)* IGV browser track showing the coverage (y-axis) of transcriptome and 5’ peaks of *dcp2*-specific genes with example of HSP23.6 (At4g25200) and transposable element gene (At2g22710) along with the expression of transcript isoforms (on right). The 5’ features identified in each sample and gene structure of encoded transcripts [UTRs (narrow rectangle), exons (broad rectangle), introns (lines connecting the exons) and arrows indicate the orientation of the gene] from IGV are displayed. *IGV*, Integrated genome viewer; *NT*, No treatment; *T*, DCP2-treatment; *TPM*, Transcripts per million; *WT*, wild type.

Further, we confirmed that genes with *dcp2*-specific caps showed low or undetectable expression in wild type, as indicated by global expression analysis (Fig. 3B) and exemplified by the cap footprints and transcript abundance of selected genes (Fig. 3C; Supplementary Fig. S4A).

This suggests that many of these genes are normally short-lived and become detectable only when decapping is impaired. Consistent with this idea, analysis of published RNA decay data showed that *dcp2*-specific capped genes display increased stability and reduced decay rates in the decapping-deficient mutant *vcs-7* compared with wild type (Supplementary Fig. S4B)(10). In addition, we identified 275 *dcp2*-specific capped loci in the intergenic regions (Supplementary Fig. S5; Supplementary Table S9). These loci represent previously unannotated short transcripts that were detected exclusively in *dcp2* mutants.

Inspection of 5’ caps localized near annotated TSSs revealed that *dcp2* plants exhibit a greater number of genes with two or more alternative 5’ features compared with the wild type (379 vs. 150; Supplementary Fig. S6). Transcript-level analysis and inspection of cap footprints indicated that the distinct 5’-end profiles observed between the two genotypes reflect the expression of different mRNA isoforms (Supplementary Fig. S6), suggesting that *dcp2*-specific alternatively capped transcripts may be less stable and less abundant. Lastly, the genes with wild type-specific caps (1,397) showed undetectable expression in *dcp2*, which also included 97 capped loci from the intergenic regions and are exemplified by the cap footprints of selected genes (Supplementary Fig. S7, Supplementary Table S8).

### Cytosolic decay substrates are processed via canonical decapping

Decapping-dependent decay is a primary pathway for the clearance of co-translationally and cytosolic decaying mRNAs in Arabidopsis (12, 63, 64, 69, 70). A recent study decoupled co-translational decay from cytosolic decay by engineering an XRN4ΔCTRD line that preserves cytosolic XRN4 activity but abolishes co-translational decay (64). We utilized published degradome datasets describing XRN4-dependent co-translational and cytosolic decay footprints and compare them with our data obtained in *dcp2* mutants. Among the total 12,306 genes expressed in the degradome dataset, 11,441 were detected in our datasets (transcript per million (TPM) ≥ 1) and showed an elevated expression in *dcp2* compared with wild type for both co-translational and cytosolic decay targets (Fig. 4A). Many genes from both groups were detected as capped in both wild type and *dcp2*. Moreover, the 5’ cap enrichment index showed higher accumulation of cap signal in *dcp2* compared with wild type (Fig. 4B). This accumulation can likely be attributed to an impaired decapping step, positioning DCP2 upstream of XRN4 within 5’-3’ decay pathways.

**Figure 4:**
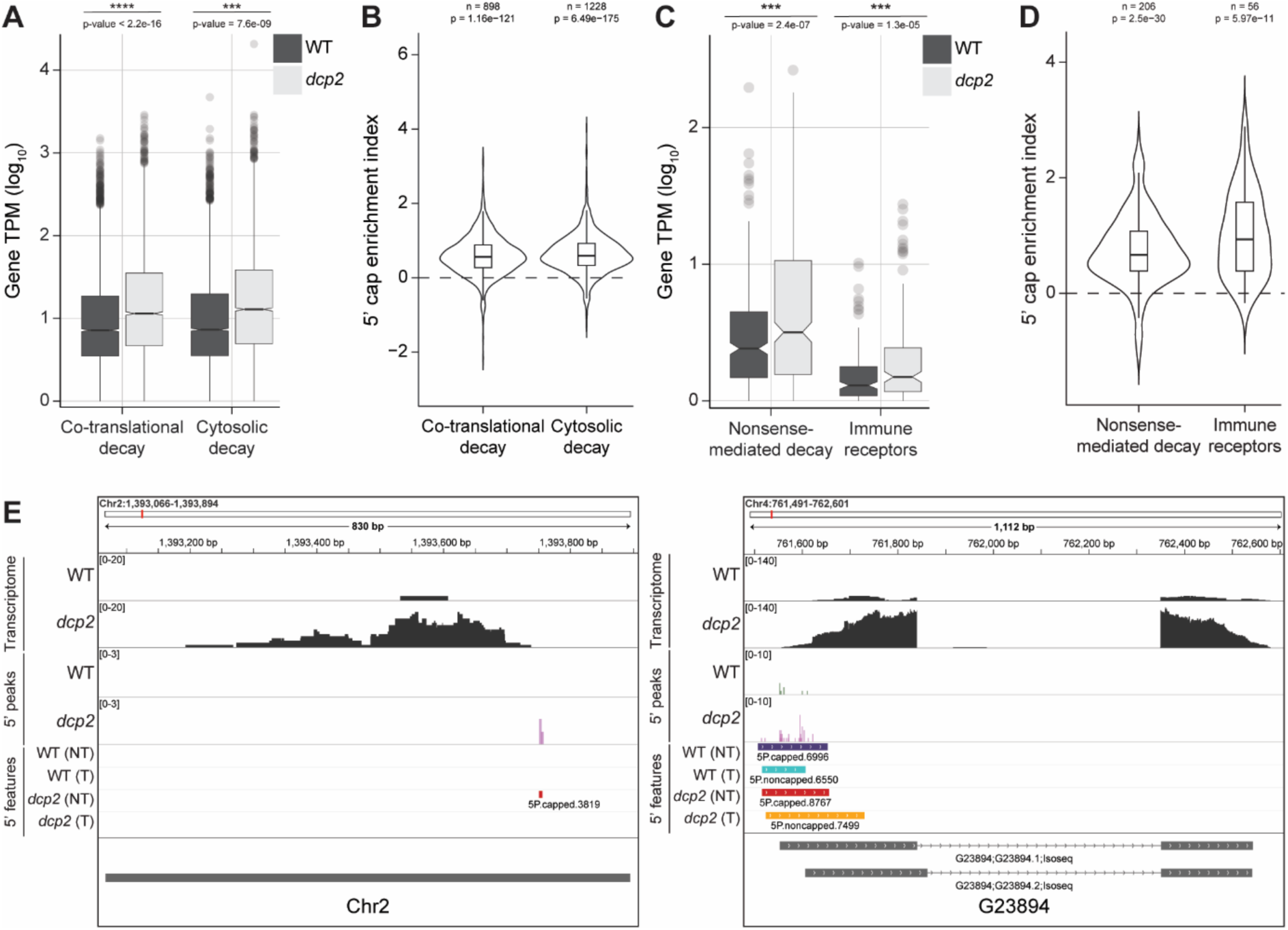
Cytosolic decay substrates are processed via canonical decapping. *(A)* Boxplot showing average gene expression (log_10_TPM) of co-translational and cytosolic decay targets that are expressed in our dataset. *(B)* Violin plot showing the distribution of 5’ cap enrichment index (log_2_) for genes undergoing co-translational or cytosolic decay target groups. Statistical significance was assessed using paired t-tests, with p-values shown. *(C)* Boxplot showing average gene expression (log_10_TPM) of NMD and immune receptors targets that are expressed in our dataset. *(D)* Violin plot showing the distribution of 5’ cap enrichment index (log_2_) for genes undergoing NMD decay. Statistical significance was assessed using paired t-tests, with p-values shown. *(E)* IGV browser track showing the coverage (y-axis) of transcriptome and 5’ peaks of selected NMD targets with example of a previously unannotated loci (left) and previously unannotated gene (G23894; on right). The 5’ features identified in each sample and the gene structure of encoded transcripts [UTRs (narrow rectangle), exons (broad rectangle), introns (lines connecting the exons) and arrows indicate the orientation of the gene] from IGV are displayed. *n*, number of genes analyzed per group; *IGV*, Integrated genome viewer; *NT*, No treatment; *T*, DCP2-treatment; *TPM*, Transcripts per million; *WT*, wild type.

Decapping-null seedlings phenotypically resemble NMD-deficient (*upf1*) mutants (Supplementary Fig. S1A) (71, 72). Genome-wide profiling of *dcp2* also revealed induction of defense and stress responses (Fig. 1C), like NMD-deficient mutants, which prompted further investigation. In plants, Enhanced Disease Susceptibility 1 (EDS1) acts as a central signaling hub during pathogen attack and coordinates other immune components and pathways to ensure a balanced defense response (73–75). Because suppressing immune signaling rescues the growth defects of NMD mutants (71, 76), we tested whether reducing immune signaling via EDS1 inactivation could alleviate decapping-related phenotypes. However, *vcs6 eds1* double mutants showed no growth rescue and displayed severe defects similar to decapping single mutants, indicating that seedling lethality is not caused solely by elevated immune responses (Supplementary Fig. S8).

Previous studies suggest that NMD often triggers RNA decay through DCP2-mediated decapping and that NMD and decapping pathways can share target transcripts (11, 12, 30, 77–81). To validate this, we used our previous dataset, where we identified 333 genes as high-confident targets of NMD along with 172 immune receptor genes, many of which are also processed by NMD (71). We examined these NMD targets in our decapping dataset.

Of the 333 high-confidence NMD targets and 172 immune receptor genes, most were detected and showed increased expression in *dcp2* mutants compared with wild type, suggesting substrate overlap between NMD and decapping pathways (Fig. 4C). Notably, the 5’ cap enrichment index of these decay target groups also showed higher accumulation of cap signal in *dcp2* compared with wild type (Fig. 4D). In addition to annotated NMD substrates, we identified several previously uncharacterized transcript isoforms and unannotated loci that are specifically expressed in *upf1* mutants and are also capped and exclusively present in *dcp2* (Fig. 4E). These observations support the conclusion that, in *dcp2*, rapidly degrading NMD targets are stabilized due to impaired decapping (Fig. 4C-D), leading to their accumulation. However, while decapping and NMD coregulate an overlapping subset of transcripts, *dcp2* seedling lethality does not appear to stem exclusively from misexpression of NMD-regulated immune receptors, but rather from a broader disruption of mRNA stability.

## DISCUSSION

The 5’ cap plays a central role in determining the stability and fate of an RNA molecule (82). In plants, bulk of cytoplasmic RNA decay is mediated by DCP2-dependent decapping followed by a 5’-3’ exonucleolytic decay pathway (10, 64). This study provides a genome-scale, nucleotide-resolved view of DCP2-dependent 5’ caps in *A. thaliana* seedlings and positions canonical decapping at the center of multiple cytoplasmic RNA turnover pathways. By combining an *in vitro* decapping with nanoPARE-based 5’-end profiling, we defined positions of 5’ mRNA caps and determined the capping status of mRNAs. The *in vitro* DCP2 treatment-dependent reduction in uuG frequency supports the specificity of the assay for bona fine 5’ caps.

Our analysis reveals a differential distribution of 5’ caps between wild type and *dcp2*, which reflects changes in the presence of transcript isoforms as well as alternate TSS between the genotypes. Alternative TSS usage is particularly important for modulating post-transcriptional regulation (67, 68). Changes in transcript abundance may reflect secondary effects on mRNA production in response to deficient decapping, as well as increased stability of rapidly degraded mRNAs. Indeed, in *dcp2* mutants we noted a prominent increase in 5‘ caps and associated transcripts, including low-abundant mRNAs and transcripts originating from previously unannotated loci. Many of these capped RNAs are barely detected in wild type, indicating that they are normally short-lived and efficiently eliminated by decapping-mediated turnover. Their stabilization in *dcp2* therefore reflects a direct consequence of impaired decapping and highlights its role in RNA homeostasis.

In the cytoplasm, once decapped, transcripts become substrates for 5’-3’ exonucleolytic degradation by the XRN4 enzyme, thereby strongly influencing overall mRNA turnover (62–66, 70). Transcripts previously classified as XRN4 substrates accumulate as capped RNAs in *dcp2* mutants and display increased cap enrichment, consistent with a block at decapping step (63, 64, 69). These findings support a model in which decapping is a prerequisite for both co-translational and cytosolic 5’-3’ decay pathways *in vivo*. In addition, our data revealed a partial overlap between canonical decapping and nonsense-mediated decay (NMD). A substantial fraction of high-confidence NMD targets, including previously unannotated unstable targets, show increased expression and cap enrichment in *dcp2*, indicating that a subset of NMD substrates relies on canonical decapping for efficient clearance (11, 71, 78).

In summary, this study provides a resource on 5’ cap distribution in Arabidopsis and establishes DCP2 as a central determinant of cap-defined RNA fate *in vivo*. Canonical decapping restricts the accumulation of unstable and cryptic transcripts, interfaces with co-translational, cytosolic, and NMD-associated decay pathways, and is essential for transcriptome integrity during early development. The cap footprint signatures defined here provide a framework for dissecting isoform-specific RNA turnover and the integration of decapping with parallel RNA surveillance pathways under developmental and stress conditions.

## ACKNOWLEDGEMENTS

We acknowledge the CF Genomics and the CF Bioinformatics supported by the NCMG research infrastructure (LM2023067 funded by MEYS CR) for their support with obtaining scientific data presented in this paper. Plant Sciences Core Facility of CEITEC Masaryk University is gratefully acknowledged for the obtaining of the scientific data presented in this study. We are thankful to the Czech Science Foundation (23-07969X) and the Ministry of Education, Youth, and Sports of the Czech Republic for the funding from the project TowArds Next GENeration Crops, reg. no. CZ.02.01.01/00/22_008/0004581 of the ERDF Programme Johannes Amos Comenius.

## AUTHOR CONTRIBUTIONS

K.R. and N.S. conceived the ideas and designed the experiments. N.S. performed the experiments and data analysis. M.A.S. and V.K.R contributed to bioinformatic analyses and data visualization. N.S., V.K.R and K.R. interpreted the results. N.S. drafted the manuscript with input from K.R. K.R. and M.D.N. supervised the project. K.R. secured the funding. All authors reviewed and approved the final manuscript.

## SUPPLEMENTARY DATA

Supplementary Data are available online.

## CONFLICT OF INTEREST

The authors declare that they have no direct or indirect competing financial or non-financial interests.

## FUNDING

This work was funded by the Czech Science Foundation (23-07969X) and by the Ministry of Education, Youth, and Sports of the Czech Republic from the project TowArds Next GENeration Crops, reg. no. CZ.02.01.01/00/22_008/0004581 of the ERDF Programme Johannes Amos Comenius.

## DATA AVAILABILITY

All the sequencing data generated in this study have been deposited in the Gene Expression Omnibus (GEO) under accession number GSE315704.

## Notes

### Competing Interest Statement

The authors have declared no competing interest.

