## Supplementary Fig. S1-S8 for "Genome-wide mapping of DCP2-dependent 5′ cap footprints in *Arabidopsis thaliana*"

### Table of contents

#### Supplementary Figures


| *Figure* | *Page* |
|---|---|
| Supplementary Figure S1: Decapping-deficient mutants in Arabidopsis. | 2 |
| Supplementary Figure S2: The direct assay method to map canonical 5′ caps. | 3 |
| Supplementary Figure S3: Changes in whole transcriptome abundance and cap enrichment. | 4 |
| Supplementary Figure S4: Examples of *dcp2*-specific capped genes. | 5 |
| Supplementary Figure S5: Examples of *dcp2*-specific capped genes originating from previously unannotated genomic loci. | 6 |
| Supplementary Figure S6: Examples of multiple 5′ caps in *dcp2* mutants along with their corresponding transcript isoforms. | 7 |
| Supplementary Figure S7: Examples of genes uniquely capped in wild type. | 8 |
| Supplementary Figure S8: The impact of EDS1 on decapping-deficient mutants in Arabidopsis. | 9 |


**Supplementary Tables** (available separately in Supplementary_Tables.xlsx)


| *Table* | *Worksheet* |
|---|---|
| Supplementary Table S1: Oligonucleotides used in present study | Table_S1 |
| Supplementary Table S2: Summary of NGS samples used in present study | Table_S2 |
| Supplementary Table S3: Unified 5' ends identified in no treatment samples by EndClass analysis | Table_S3 |
| Supplementary Table S4: 5' ends identified in wild type (No treatment) | Table_S4 |
| Supplementary Table S5: 5' ends identified in *dcp2* mutant (No treatment) | Table_S5 |
| Supplementary Table S6: 5' ends identified in wild type after *in vitro* DCP2 treatment (DCP2-resistant 5' ends in wild type) | Table_S6 |
| Supplementary Table S7: 5' ends identified in *dcp2* after *in vitro* DCP2 treatment (DCP2-resistant 5' ends in *dcp2*) | Table_S7 |
| Supplementary Table S8: High-confident 5' caps identified in wild type | Table_S8 |
| Supplementary Table S9: High-confident 5' caps identified in *dcp2* | Table_S9 |

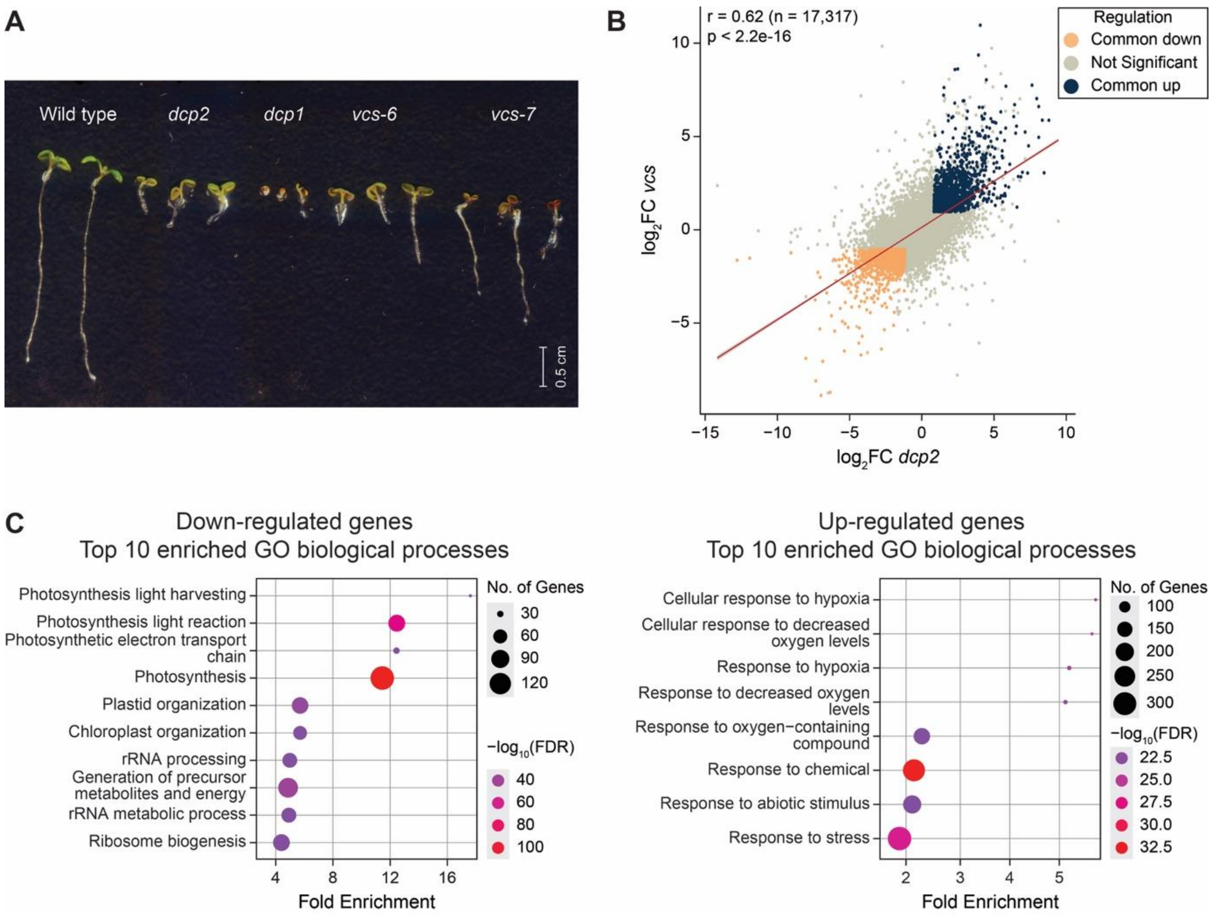


**Supplementary Figure S1:** Decapping-deficient mutants in Arabidopsis. *(A)* Phenotypes of six-day-old seedlings of wild type and decapping-deficient null mutants (*dcp2*, *dcp1*, *vcs-6*, *vcs-7*). *(B)* Scatter plot of $\log_2$ fold change in gene expression (versus respective wild type samples) comparing transcriptomes of decapping-deficient mutants (*dcp2* and *vcs-7*). Each dot represents the $\log_2$FC of a gene's expression in either *dcp2* (x-axis) or *vcs-7* (y-axis) relative to wild type. The Pearson correlation coefficient ($r = 0.62$, $n = 17{,}317$, $p < 2.2e-16$) is displayed. *(C)* Gene Ontology enrichment analysis with dot plots showing top 10 enriched GO biological processes among shared up- and down-regulated genes in *dcp2* and *vcs-7* compared to their respective wild type (adjusted p-value < 0.05). The most to least significantly enriched categories are colored as red to purple, and the size of circle correspond to the number of genes in each category. *FC*, fold change; *n*, total number of genes; *GO*, gene ontology.

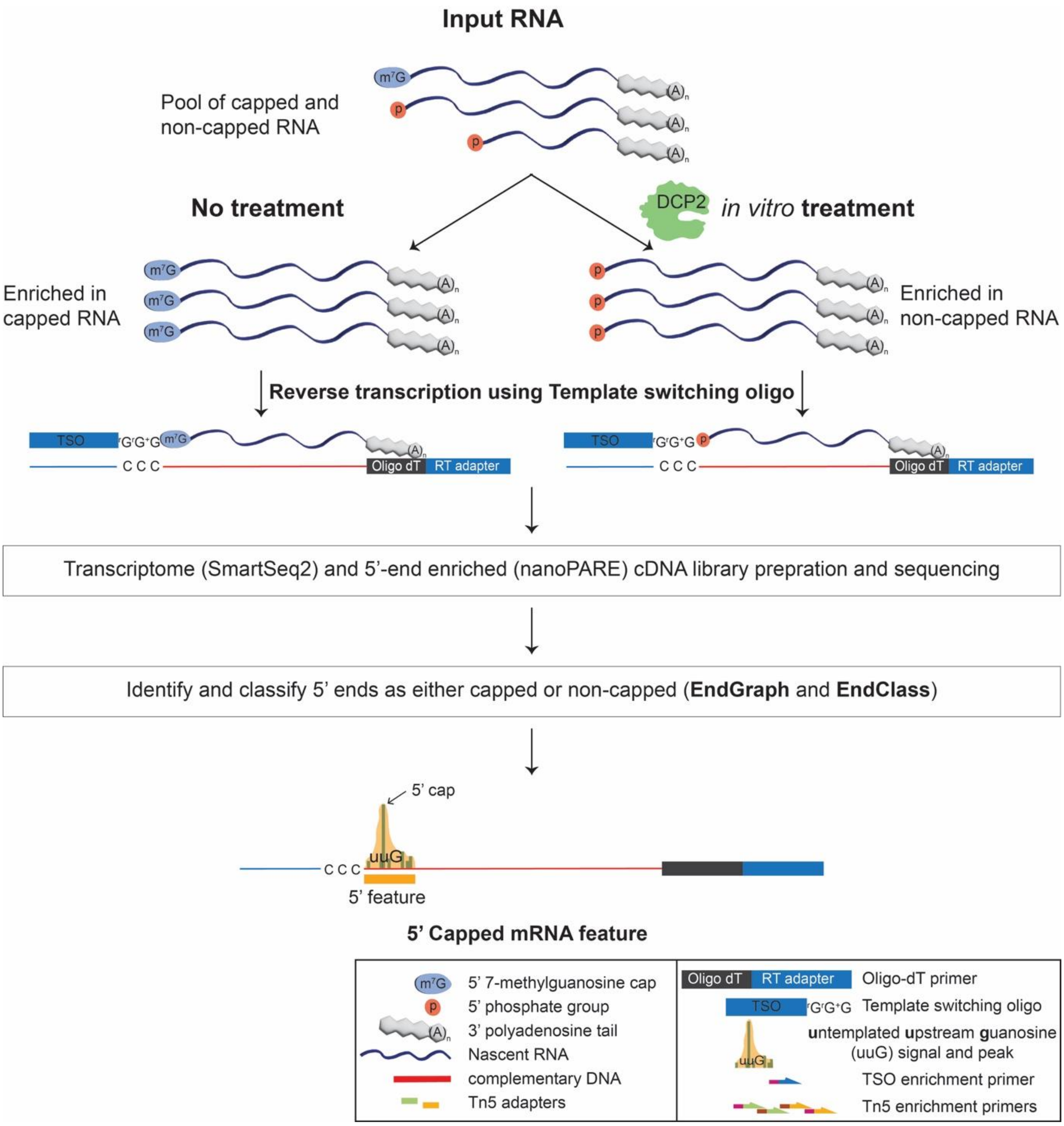


**Supplementary Figure S2:** The direct assay method to map canonical 5′ caps. Schematic depiction of direct assay method involving *in vitro* decapping treatment of total RNA that hydrolyzes 5′ $m^7G$ cap, thus increasing the proportion of non-capped RNA. Subsequently, SmartSeq2 and low input 5′-end enriched (Parallel analysis of RNA ends (nanoPARE)) cDNA libraries were prepared from polyA-selected RNA of both no treatment and *in vitro* treated samples. Later, the nanoPARE data analysis pipeline identifies and classifies the 5′ features as either capped or non-capped based on untemplated upstream guanosine peak signal. The direct assay methodology aids in identification of high-confidence 5′ caps at single nucleotide resolution by utilizing the *in vitro* treated sample in combination with the mutant. *$m^7G$*, methylguanosine cap; *uuG*, untemplated upstream guanosine.

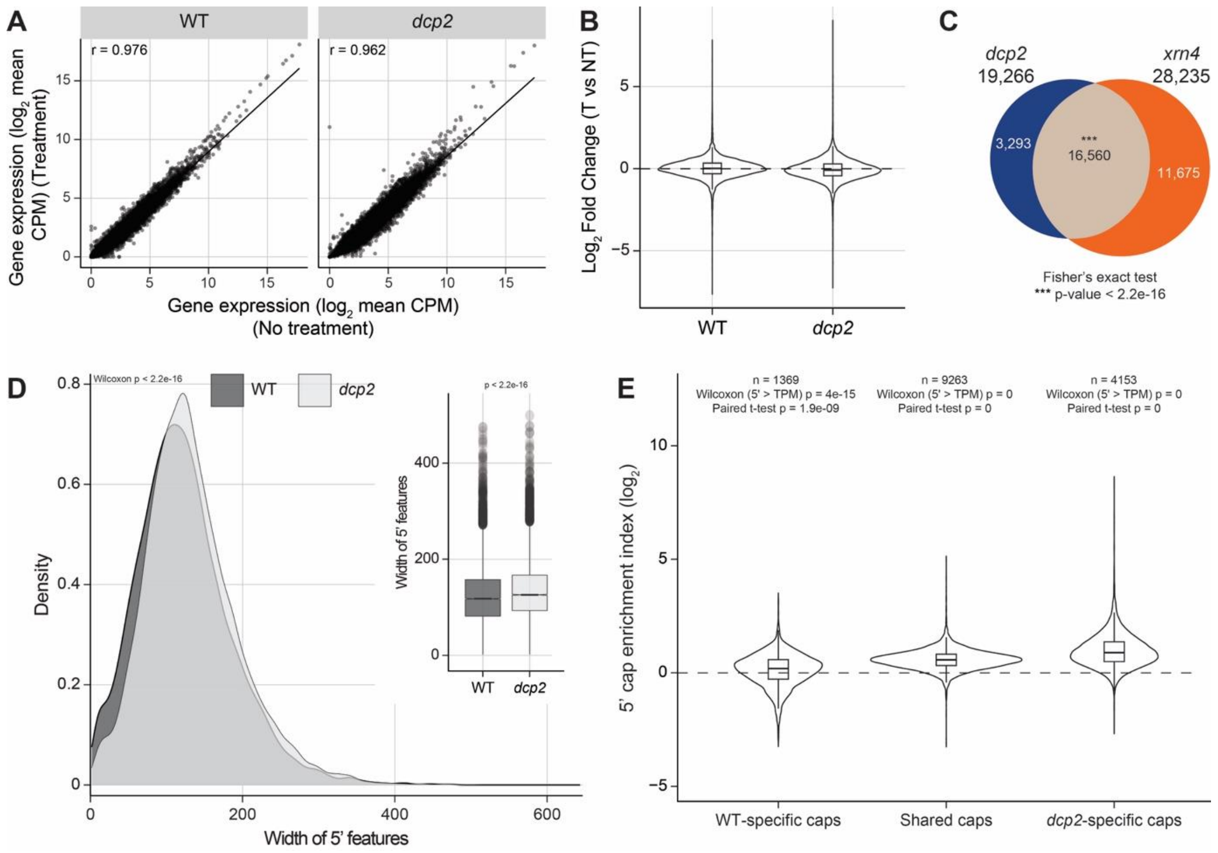


**Supplementary Figure S3:** Changes in whole transcriptome abundance and cap enrichment. *(A)* Scatter plots of gene abundance (mean $log_2$ (CPM)) between conditions (treatment vs no treatment) and Pearson correlation coefficient per genotype with statistical significance based on student's t-test. The dashed line is linear regression fit. *(B)* Violin plots of gene-wise fold-change distributions between no treatment versus treated samples in both wild type and *dcp2* mutant. *(C)* Venn diagram showing the comparison between 5′ features identified in *dcp2* (present study) and *xrn4* (Schon et al., 2018). The overlapping region (brown) are the 5′ features shared between the samples while the non-overlapping regions depict the 5′ features unique to either *dcp2* (blue) or *xrn4* (orange). The asterisk mark indicates the significance of overlap between the groups, determined by the Fisher's exact test. *(D)* Density and boxplot (inset) for width analysis of high-confidence 5′ features. *(E)* Violin plot showing the distribution of 5′ cap enrichment index ($log_2$) for capped genes classified as wild type-specific, shared or *dcp2*-specific. Statistical significance was assessed using paired Wilcoxon rank-sum test and paired t-tests, with p-values shown. *CPM*, counts per million; *r*, Pearson correlation coefficient; *n*, number of genes analyzed per group; *NT*, No treatment; *T*, DCP2-treatment; *WT*, wild type.

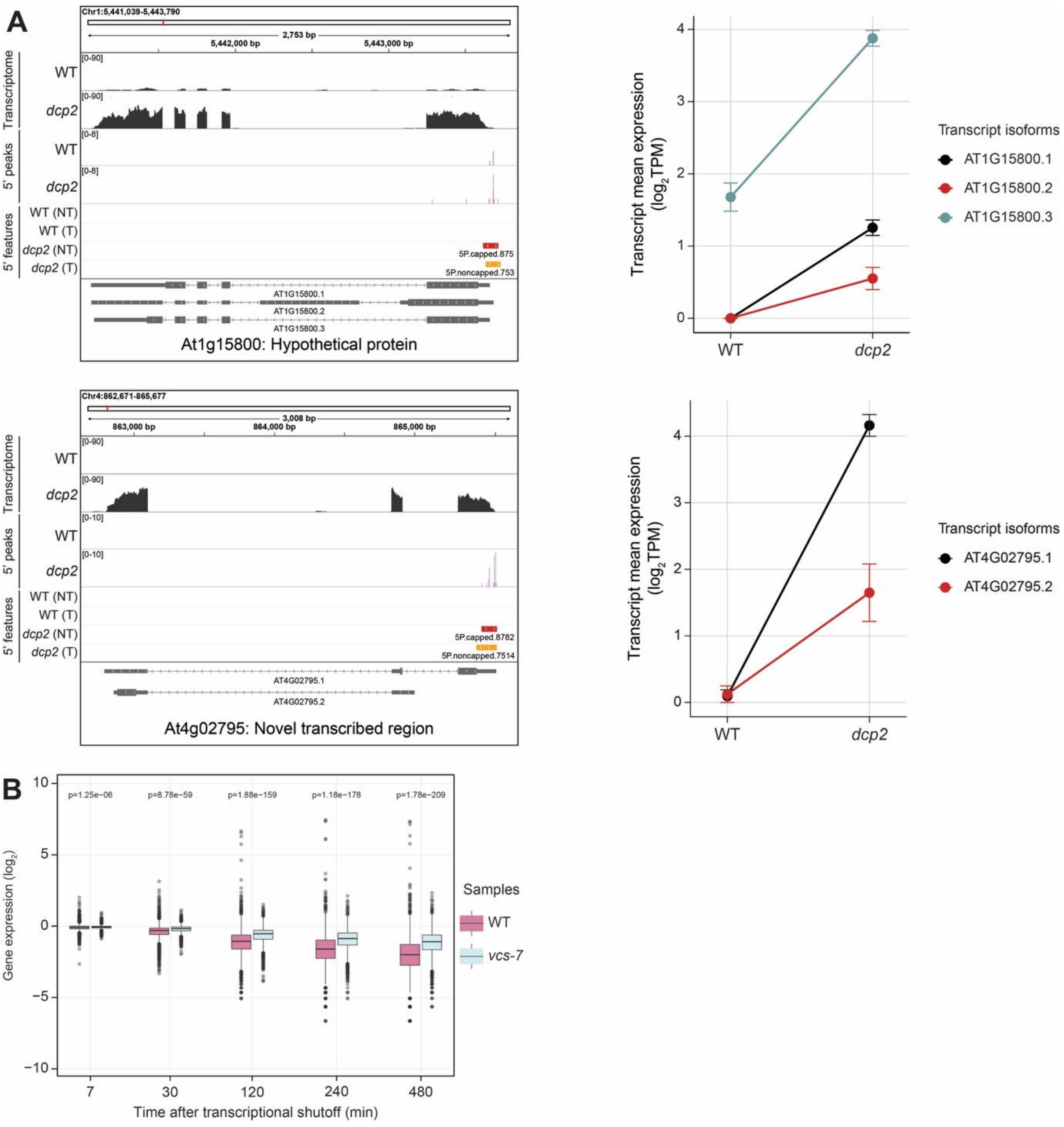


**Supplementary Figure S4:** Examples of *dcp2*-specific capped genes. *(A)* IGV browser tracks showing the coverage (y-axis) of transcriptome and 5′ peaks of *dcp2*-specific capped genes along with selected transcript isoforms expression (on right). The 5′ features identified in each sample and gene structure of encoded transcripts [UTRs (narrow rectangle), exons (broad rectangle), introns (lines connecting the exons) from IGV are displayed. *(B)* RNA decay kinetics of *dcp2*-specific capped genes in wild type (pink) and *vcs-7* (blue). Boxplots show the distribution of relative gene expression ($log_2$) across time points following transcriptional shutoff (Sorenson et al., 2018). The p-values above each time point represent the statistical significance of expression differences between wild type and *vcs-7* (two-sided Student's t-test, Benjamini-Hochberg adjusted). *IGV*, Integrated genome viewer; *NT*, No treatment; *T*, DCP2-treatment; *WT*, wild type.

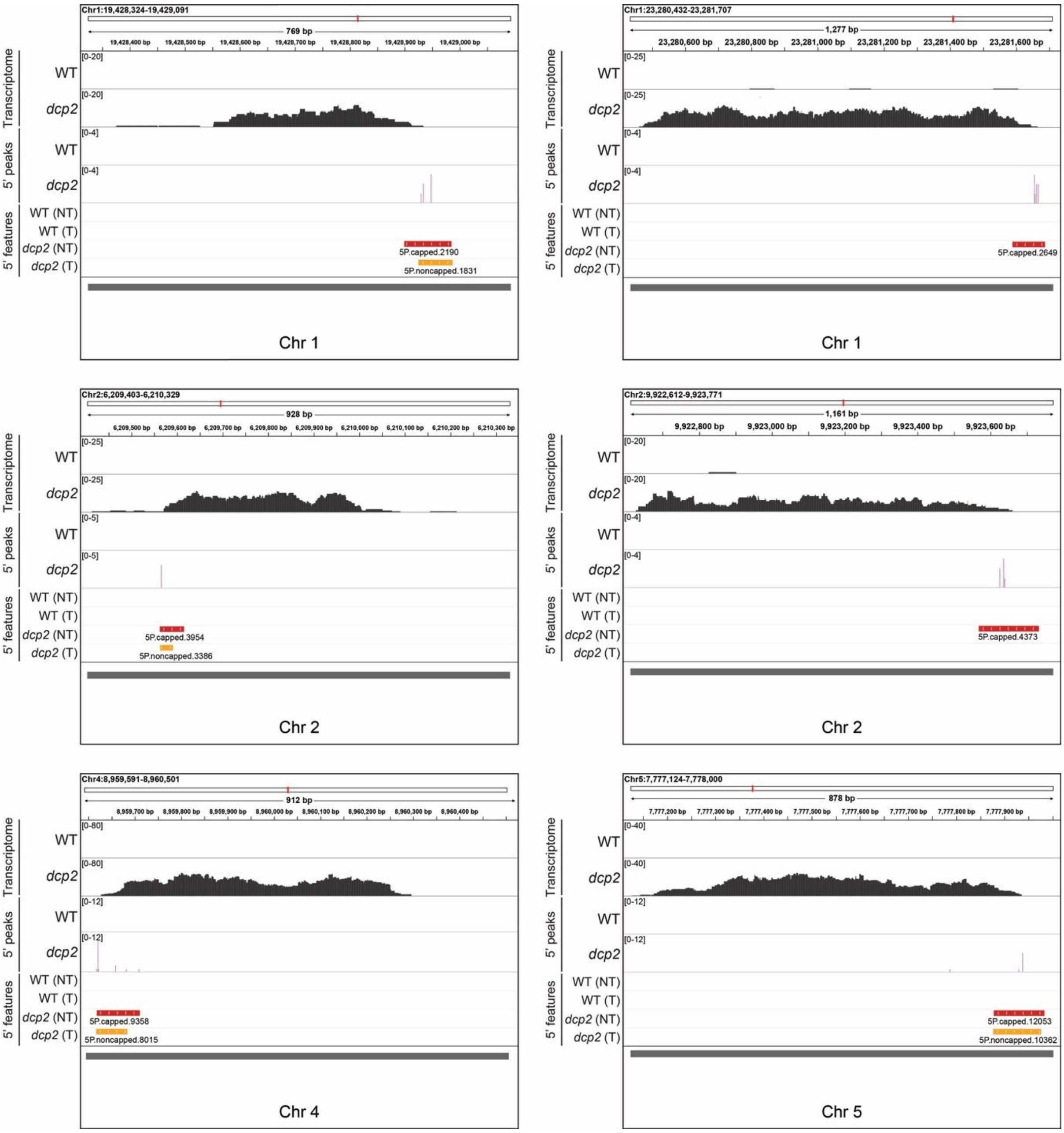


**Supplementary Figure S5:** Examples of *dcp2*-specific capped genes originating from previously unannotated genomic loci. IGV browser tracks showing the coverage (y-axis) of transcriptome and 5′ peaks of *dcp2*-specific capped loci originating from the intergenic regions of genome. The 5′ features identified in each sample and the gene structure of encoded transcripts [UTRs (narrow rectangle), exons (broad rectangle), introns (lines connecting the exons) and arrows indicate the orientation of the gene] from IGV are displayed. *IGV*, Integrated genome viewer; *NT*, No treatment; *T*, DCP2-treatment; *WT*, wild type.

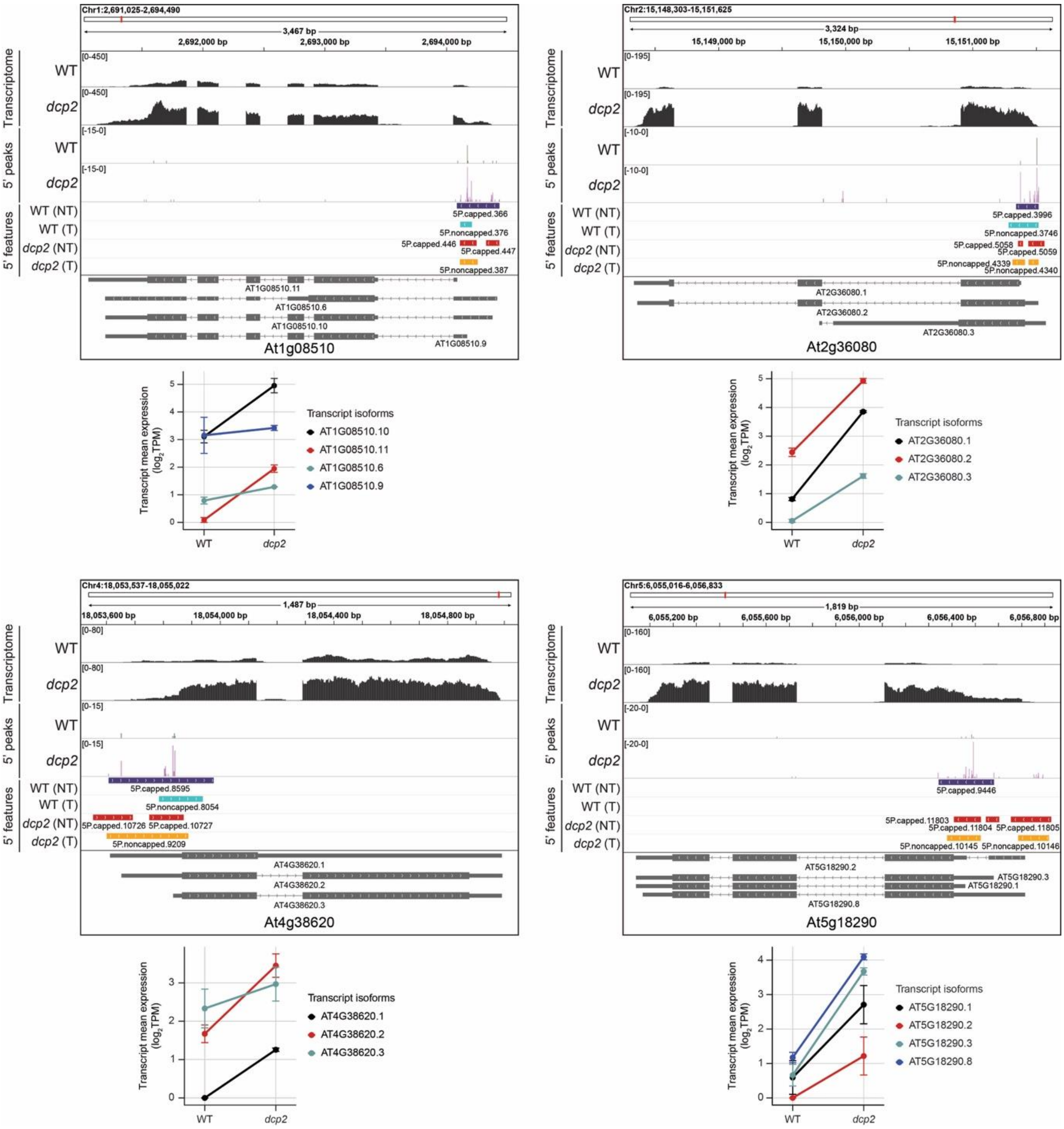


**Supplementary Figure S6:** Examples of multiple 5′ caps in *dcp2* mutants along with their corresponding transcript isoforms. IGV browser tracks showing the coverage (y-axis) of transcriptome and 5′ peaks of genes with more than one 5′ cap being assigned to one gene in *dcp2* along with the expression of transcript isoforms (bottom of each IGV track). The 5′ features identified in each sample and the gene structure of encoded transcripts [UTRs (narrow rectangle), exons (broad rectangle), introns (lines connecting the exons) and arrows indicate the orientation of the gene] from IGV are displayed. *IGV*, Integrated genome viewer; *NT*, No treatment; *T*, DCP2-treatment; *WT*, wild type.

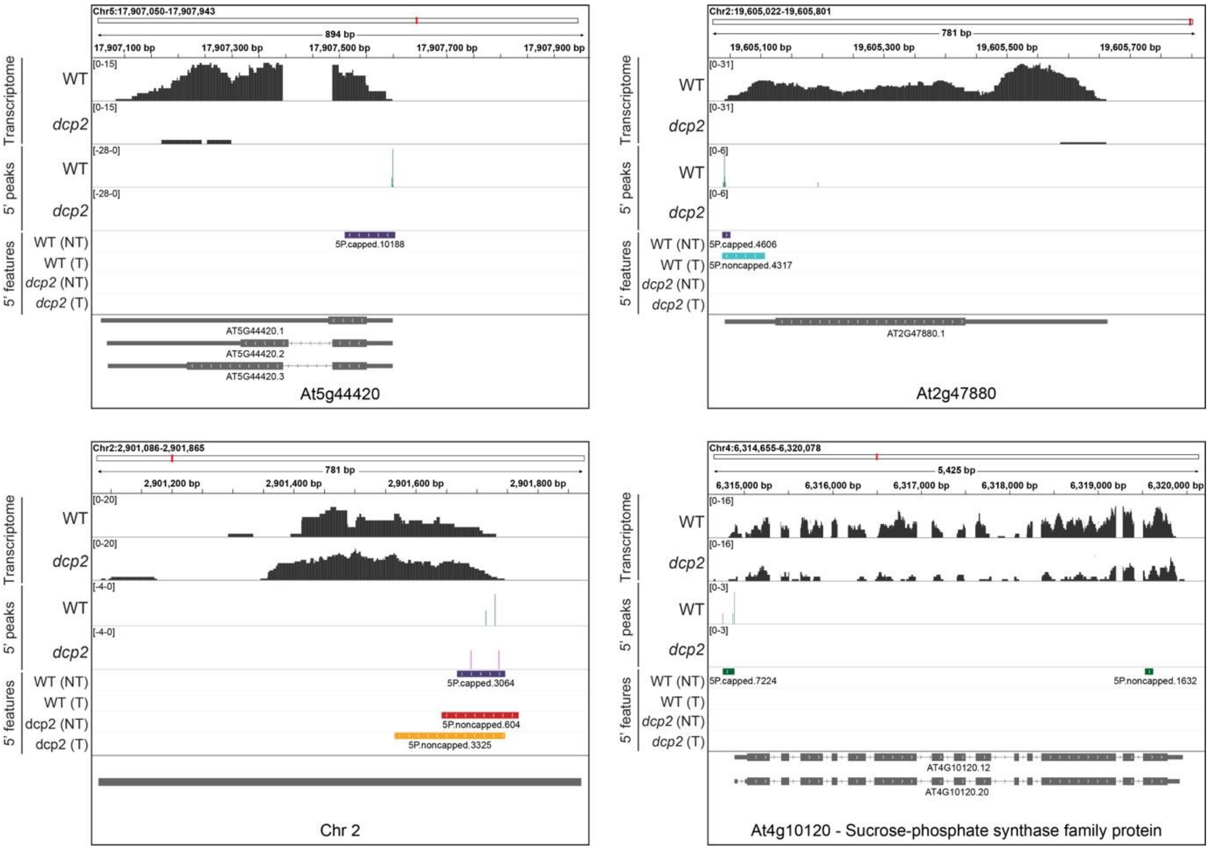


**Supplementary Figure S7:** Examples of genes uniquely capped in wild type. IGV browser tracks showing the coverage (y-axis) of transcriptome and 5′ peaks of wild type-specific capped genes. The 5′ features identified in each sample and the gene structure of encoded transcripts [UTRs (narrow rectangle), exons (broad rectangle), introns (lines connecting the exons) and arrows indicate the orientation of the gene] from IGV are displayed. *IGV*, Integrated genome viewer; *NT*, No treatment; *T*, DCP2-treatment; *WT*, wild type.

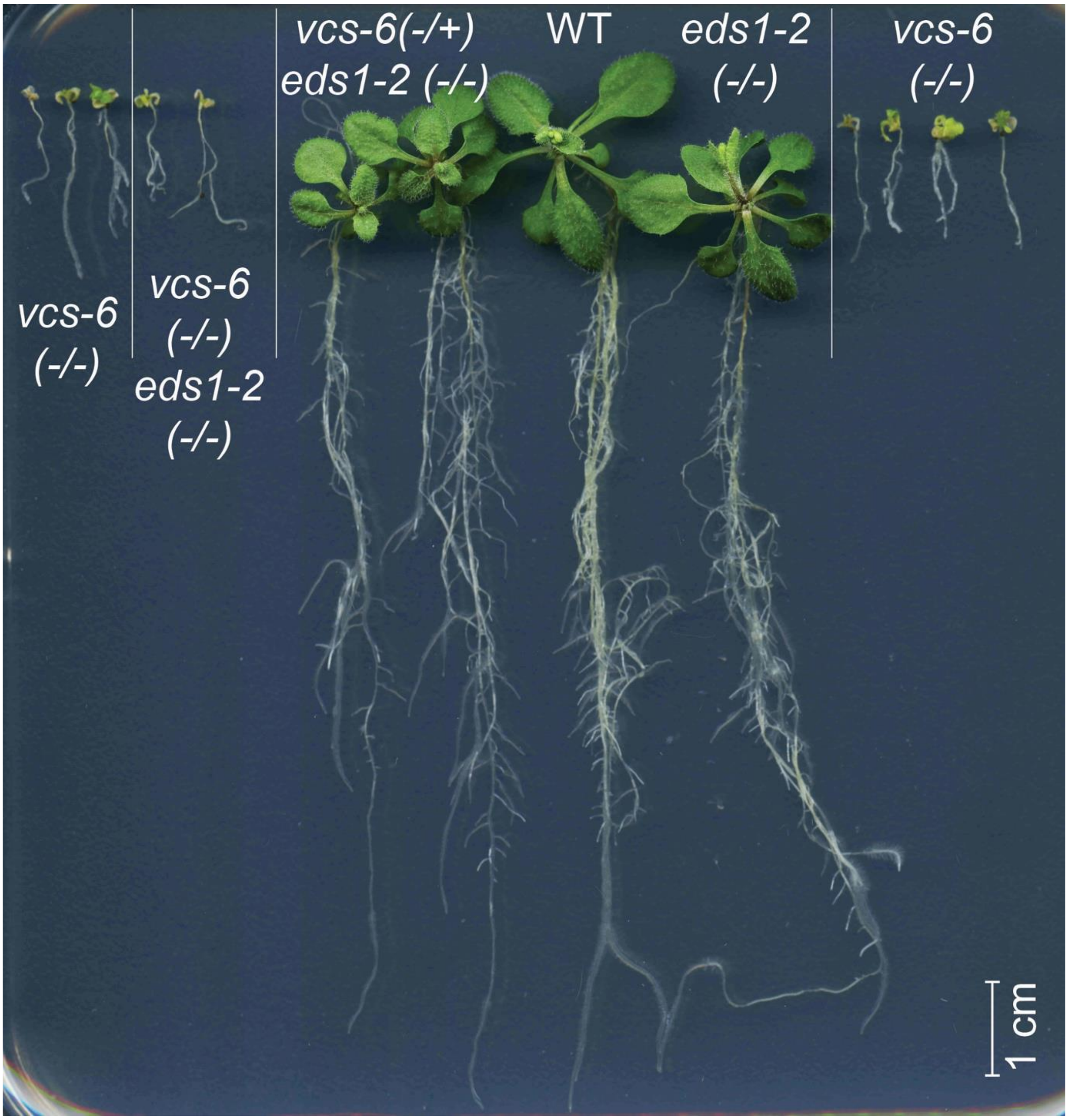


**Supplementary Figure S8:** The impact of EDS1 on decapping-deficient mutants in Arabidopsis. Phenotypes of 21-day old seedlings of wild type, immune-supressed (*eds1*) mutant and decapping-deficient mutants (*vcs-6*), with a no rescue observed in *vcs6eds1* double mutants. *WT*, wild type.
